# Functional basis for calmodulation of the TRPV5 calcium channel

**DOI:** 10.1101/2021.02.16.431366

**Authors:** Sara R Roig, Niky Thijssen, Merijn van Erp, Jack Fransen, Joost G Hoenderop, Jenny van der Wijst

## Abstract

Within the transient receptor potential (TRP) superfamily of ion channels, TRPV5 is a highly Ca^2+^-selective channel important for active reabsorption of Ca^2+^ in the kidney. Its channel activity is controlled by a negative feedback mechanism involving calmodulin (CaM) binding. Combining advanced microscopy techniques and biochemical assays, this study characterized the dynamic bilobal CaM regulation and binding stoichiometry. We demonstrate for the first time that functional (full-length) TRPV5 interacts with CaM in the absence of Ca^2+^, and this interaction is intensified at increasing Ca^2+^ concentrations sensed by the CaM C-lobe that achieves channel pore blocking. Channel inactivation occurs without CaM N-lobe calcification. Moreover, we reveal a 1:2 stoichiometry of TRPV5:CaM binding by implementing *single molecule photobleaching counting*, a technique with great potential for studying TRP channel regulation. In conclusion, our study proposes a new model for CaM-dependent regulation – *calmodulation* – of the Ca^2+^-selective TRPV5 that involves apoCaM interaction and lobe-specific actions.

## Introduction

Transient receptor potential (TRP) channels are one of the largest classes of ion channels and are widely expressed throughout the animal kingdom (1). Since their discovery, they have emerged as key players in human physiology and were found to be associated with various diseases, such as cancer, skeletal abnormalities, skin disorders, and chronic pain (2, 3). The mammalian TRP family consists of six subfamilies; canonical (TRPC), melastatin (TRPM), vanilloid (TRPV), polycystin (TRPP), ankyrin (TRPA) and mucolipin (TRPML), categorized based on sequence homology (4).

Yet, there is a striking variability in their channel properties in contrast to other families of ion channels. While most TRP channels are rather non-selective for ion permeation, TRPV5 and its close homologue TRPV6 exhibit a highly selective calcium (Ca^2+^) permeability (5, 6). This concurs with their transport function in Ca^2+^ (re)absorbing epithelia of the kidney and intestine. Functionally, the channels utilize a Ca^2+^-dependent feedback mechanism to achieve a fast channel inactivation and slow current decay (7, 8). Over the years, it has been established that (part of) this regulation occurs via interaction with the Ca^2+^-binding protein calmodulin (CaM) (9-14).

CaM regulation is well studied across voltage-gated Ca^2+^ channels (Ca_v_), where it is also referred to as ‘calmodulation’ (15). CaM consists of amino (N)-terminal and carboxy (C)- terminal lobes, each containing two EF hand pairs capable of binding Ca^2+^, that are joined by a flexible central linker. CaM can customize channel regulation by binding Ca^2+^ to either its N- or C-lobe and evoke lobe-specific channel modulation. Moreover, it is known to bind Ca_v_ channels in its Ca^2+^-free state (apoCaM), and can consequently transduce relevant signals upon Ca^2+^ binding (15). Hence, a prominent feedback mechanism can tune channel gating, and thus regulate Ca^2+^ influx in accordance with cytosolic Ca^2+^ signals. While CaM is known to bind various TRP channels, including TRPV5/6, surprisingly little is known about such ‘TRP calmodulation’ regarding the dynamics of the process and stoichiometry of binding.

Importantly, technological advances in single particle cryo-electron microscopy (cryo-EM) have demonstrated the transformative power for generating high resolution structures of TRP channels. Three independent groups, including ours, have provided detailed structural insight into TRPV5 and TRPV6 in complex with CaM (16-18). All structures were resolved upon purifying the complex in buffers containing a high Ca^2+^ concentration ([Ca^2+^]; 5 mM). This universally demonstrated binding of the CaM N- and C-lobes to respectively proximal and distal C-terminal regions of the channel (16-18). Moreover, a general blockade mechanism was shown by the side chain of K115 of the CaM C-lobe that protrudes into the channel pore. However, one of the important differences is the complex stoichiometry. Singh *et al.* and Hughes *et al.* showed that both the TRPV6-CaM and TRPV5-CaM, respectively, exhibit 1:1 stoichiometry (18), while a flexible 1:2 stoichiometry of 1 tetrameric channel binding to 2 CaM molecules was postulated based on our TRPV5-CaM complex structure (16). In order to fully understand channel regulation, it is highly significant to unravel the dynamic interaction of CaM with TRPV5, including apoCaM binding, lobe-specific effects and binding stoichiometry.

Using a combination of Förster Resonance Energy Transfer (FRET)-based Fluorescence Lifetime Imaging Microscopy (FLIM; FLIM-FRET) and Fura-2 Ca^2+^ imaging, we revealed that apoCaM is pre-associated with TRPV5. The Ca^2+^-insensitive CaM mutants, where either two EF hands (CaM12 or CaM34) or all four EF hands (CaM1234) were mutated, showed that calcification of the CaM C-lobe is essential for channel inactivation, while a calcified N-lobe appears not critical for TRPV5 inhibition. In contrast to previously proposed concepts (9, 10), this suggests that the CaM N-lobe does not function as a Ca^2+^ sensor towards rearranging the TRPV5-CaM complex in an inactivated state. Moreover, our study determined a fixed 1:2 TRPV5-CaM stoichiometry by single molecule photobleaching with total internal reflection fluorescence microscopy (SMP-TIRFM), which may be highly relevant for fast channel inactivation.

## Results

### Constitutive interaction of TRPV5 and CaM

Over the last decades, several groups focused the attention on the TRPV5-CaM interaction (10-13, 19). Biochemical studies and NMR (nuclear magnetic resonance) analysis showed that TRPV5 binds to Ca^2+^-CaM (10-13). However, these studies only used two distinct options: complete absence (EGTA) or high levels of Ca^2+^, and are mainly based on truncated versions of the channel. Recent cryo-EM structures of TRPV5 in complex with CaM exposed their exact interaction interface and postulated how CaM blocks the TRPV5 channel pore in the presence of Ca^2+^ (16, 17). Yet, the dynamic interaction between TRPV5 and CaM that conducts to the channel blocking are poorly understood.

We implemented the Forster Resonance Energy Transfer technique (FRET) using Fluorescence Lifetime Imaging Microscopy (FLIM) to unravel the Ca^2+^-dependency of interaction between TRPV5 and CaM. To this end, wildtype and well-established (partly) Ca^2+^-insensitive mutants of CaM (CaM, CaM12, CaM34, CaM1234) were fused to eGFP, and co-expressed with mCherry or mCherry-tagged TRPV5 in HEK293 cells (**Figure 1A-B**). These CaM mutants are deficient in binding Ca^2+^ due to point mutations in the N-lobe EF hands (CaM12), C-lobe EF hands (CaM34) or all four EF hands (CaM1234). Expression and distribution of all CaM mutants resembles CaM wildtype (**Figure 1A**). Co-expression with TRPV5 shifted the distribution of CaM and CaM12 from a whole-cell staining to an extranuclear distribution (**Figure 1A**). Lifetime decay (τ) measurements performed in areas of co-localization showed a significantly decreased τ of eGFP in cells co-expressing TRPV5 with CaM or CaM12 (**Figure 1B**). CaM34 and CaM1234 distributed mainly in the nuclei showing high values of decay due to its concentration. Importantly, only extranuclear ROIs were analysed, to compare between mock and TRPV5 conditions as TRPV5 is not localized in the nuclei. CaM1234 depicted a modest but significant decrease of lifetime decay, once co-expressed with TRPV5. Interestingly, CaM34 does not show any changes in τ (**Figure 1B**). Thereby, CaM, CaM12 and CaM1234 have the potential to interact with TRPV5, indicating that the CaM N-lobe does not need to calcified in order to bind TRPV5.

**Figure 1.**
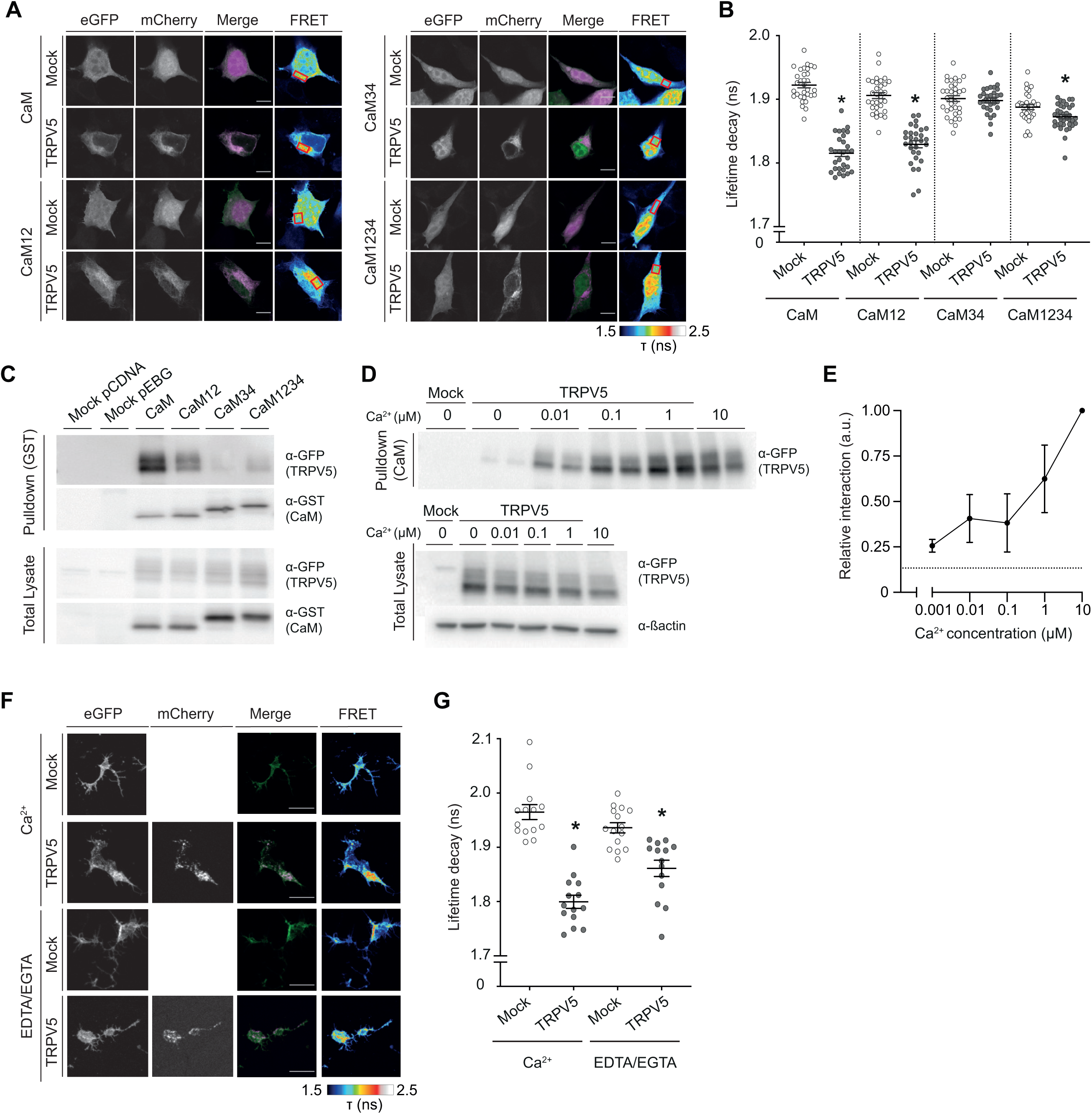
Constitutive and Ca^2+^-dependent interaction of TRPV5 and CaM. **A**. Interaction of mCherry-TRPV5 with CaM-eGFP and the Ca^2+^-insensitive mutants CaM12-eGFP, CaM34- eGFP and CaM1234-eGFP in HEK293 cells was addressed by FLIM-FRET. Representative images are shown for the donors (CaM-eGFP and indicated mutants) and acceptor (mCherry-TRPV5) expression, merged channels (donors: green, acceptor: magenta, co-localization: light pink or white) and FLIM-FRET. Example of measured ROIs represented as red boxes on the FLIM-FRET image. Bars represent 10 μm. **B**. Life time decay of CaM wildtype and mutants in the presence of mock or TRPV5, depicted as single measurements of ROIs at cytoplasmic colocalization (n=30-38 cells). * indicates p<0.05 compared to mock within each condition (ANOVA). **C**. Immunoprecipitations of HEK293 cells expressing either mock eGFP (pcDNA), mock GST (pEBG), or GST-tagged CaM wildtype and indicated mutants with eGFP-TRPV5. Representative immunoblots are shown for total lysate (bottom two lanes) and immunoprecipitated fraction of a GST pulldown (top two lanes). **D**. CaM agarose pulldown of HEK293 cells expressing either mock eGFP or eGFP-TRPV5, in the presence of increasing Ca^2+^ concentrations in the lysis buffer. Representative immunoblots are shown for the pulldown (top lane) of eGFP-TRPV5 and total lysate (bottom two lanes) of TRPV5 and ß-actin, as loading control. **E**. Semi-quantification of CaM pulldown experiments expressed as mean ± SEM (n=2-4). Intensity of TRPV5 pulldown lanes were normalized to TRPV5 total lysate. Dotted line indicates value at 0 µM Ca^2+^. **F**. FLIM-FRET analysis of mCherry-TRPV5 with CaM-eGFP in PMLs of HEK293 cells washed with either Ca^2+^-containing (2 mM) or Ca^2+^-free (2 mM EGTA, 2 mM EDTA) buffer. Representative images are shown for the donor (CaM-eGFP) and acceptor (mCherry-TRPV5) expression, merged channels (donor: green, acceptor: magenta, co-localization: light pink or white) and FLIM-FRET. Bars represent 10 μm. **G**. Life time decay of CaM in the presence of mock or TRPV5, depicted as single measurements of ROIs, for either Ca^2+^-containing or Ca^2+^-free condition (n=14-15 cells). ROIs for analysis were drawn throughout the PML. * indicates p<0.05 compared to mock within each condition (ANOVA).

We confirmed these results by co-immunoprecipitation of GFP-TRPV5 with GST-tagged CaM wildtype and mutants (**Figure 1C**). All CaM proteins demonstrated similar expression by equal pull-down with GST-sepharose resin (**Figure 1C, pulldown GST**). TRPV5 shows a clear interaction with CaM and CaM12. In line with the FLIM-FRET results, CaM1234 maintains the capability of interacting with TRPV5, albeit to lesser extent compared to CaM WT or CaM12, while CaM34 shows little-to-no interaction with TRPV5 (**Figure 1C, pulldown GST**). Of note, total lysate shows equal expression of all proteins (**Figure 1C, total lysate**).

CaM consists an N-lobe and C-lobe that each bind Ca^2+^ at different affinity (20). While the C-lobe presents a 10-fold higher affinity than the N-lobe, the kinetics is much slower (20). To understand the magnitude of Ca^2+^-dependency of CaM interaction with TRPV5, and the contribution of each lobe, a CaM agarose pull-down of TRPV5-expressing HEK293 cells was performed under different Ca^2+^ concentrations mimicking intracellular Ca^2+^ levels (0-10 µM) (**Figure 1D**). These were calculated as described in the *Methods*. Interestingly, we observed a weak interaction with TRPV5 in the absence of Ca^2+^, that was intensified upon increasing Ca^2+^ concentrations (**Figure 1D-E**). Most notably, significant interaction was already detectable at 10 nM Ca^2+^. Based on literature (20), only the C-lobe is loaded with Ca^2+^ at this specific concentration, responsible for the Ca^2+^-dependent interaction. These results point towards a pre-existing TRPV5-CaM interaction at Ca^2+^ basal concentrations.

Since we also observed interaction between TRPV5 and CaM1234, we suggest that the TRPV5-CaM complex remains even if Ca^2+^ decreases in specific subcellular areas. Yet, it might be argued that CaM1234 is structurally different to apoCaM and does not represent a physiological setting. To understand the apoCaM interaction, we performed by FLIM-FRET on PMLs. Importantly, these PMLs were prepared by unroofing the cells (**Figure 1 – figure supplement 1A**) and provided access to the intracellular compartment. Only proteins associated to the membrane remain in these preparations (21, 22). We first confirmed the presence of TRPV5-containing membranes (**Figure 1 – figure supplement 1B-C**). Following, we studied the presence of eGFP-CaM in PMLs treated with Ca^2+^-containing (2 mM) and Ca^2+^-free (2 mM EGTA, 2 mM EDTA) solutions. Despite that CaM is a cytosolic protein, CaM and TRPV5 positively co-localized in all PML conditions (**Figure 1F**). FLIM-FRET measurements proved that not only Ca^2+^-CaM, but also apoCaM interacts with TRPV5, although in a lesser extent (**Figure 1G**).

### Interaction interface of TRPV5-CaM

While the recently published cryo-EM structures of the TRPV5-CaM complex revealed that the CaM C-lobe is occluding the pore leading to channel inhibition (16, 17), no information has been provided about the dynamic bilobal regulation of TRPV5. A proximal C-terminal helix of TRPV5 was shown to bind the CaM N-lobe, while a distal C-terminal helix interacts with the CaM C-lobe (16, 17). To investigate the importance of the independent CaM lobes in binding TRPV5, we synthesized peptides of the corresponding TRPV5 helices – proximal peptide and distal peptide (**Figure 2A**). Binding of these peptides to CaM was assessed by a peptide pull-down on wildtype CaM-expressing HEK293 cells (**Figure 2B**). This demonstrated that CaM interacts with both TRPV5 peptides (**Figure 2B**). Furthermore, the peptides were used in a peptide competition assay to analyse differences in binding affinity to CaM. Addition of increasing amounts of peptide (0-100 µM) to a CaM agarose pull-down of TRPV5-expressing HEK293 cell lysates revealed decreased binding of TRPV5 to CaM-agarose by the distal peptide (**Figure 2C**). Addition of the proximal peptide resulted in significant less changes, suggesting a specific competition of the distal peptide (**Figure 2C**). Non-linear fitting of the curves to the log concentration revealed a sigmoidal correlation (R^2^= 0.942) for the distal peptide (**Figure 2E**), while no correlation was observed (R^2^= 0.188) for the proximal peptide (**Figure 2D**). Based on the fitted formula describing the graph, we calculated an IC_50_ of 5.2 µM of the competing distal peptide.

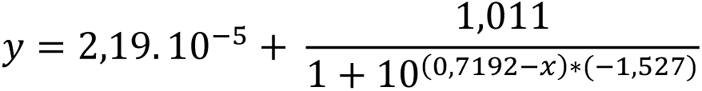

**Figure 2.**
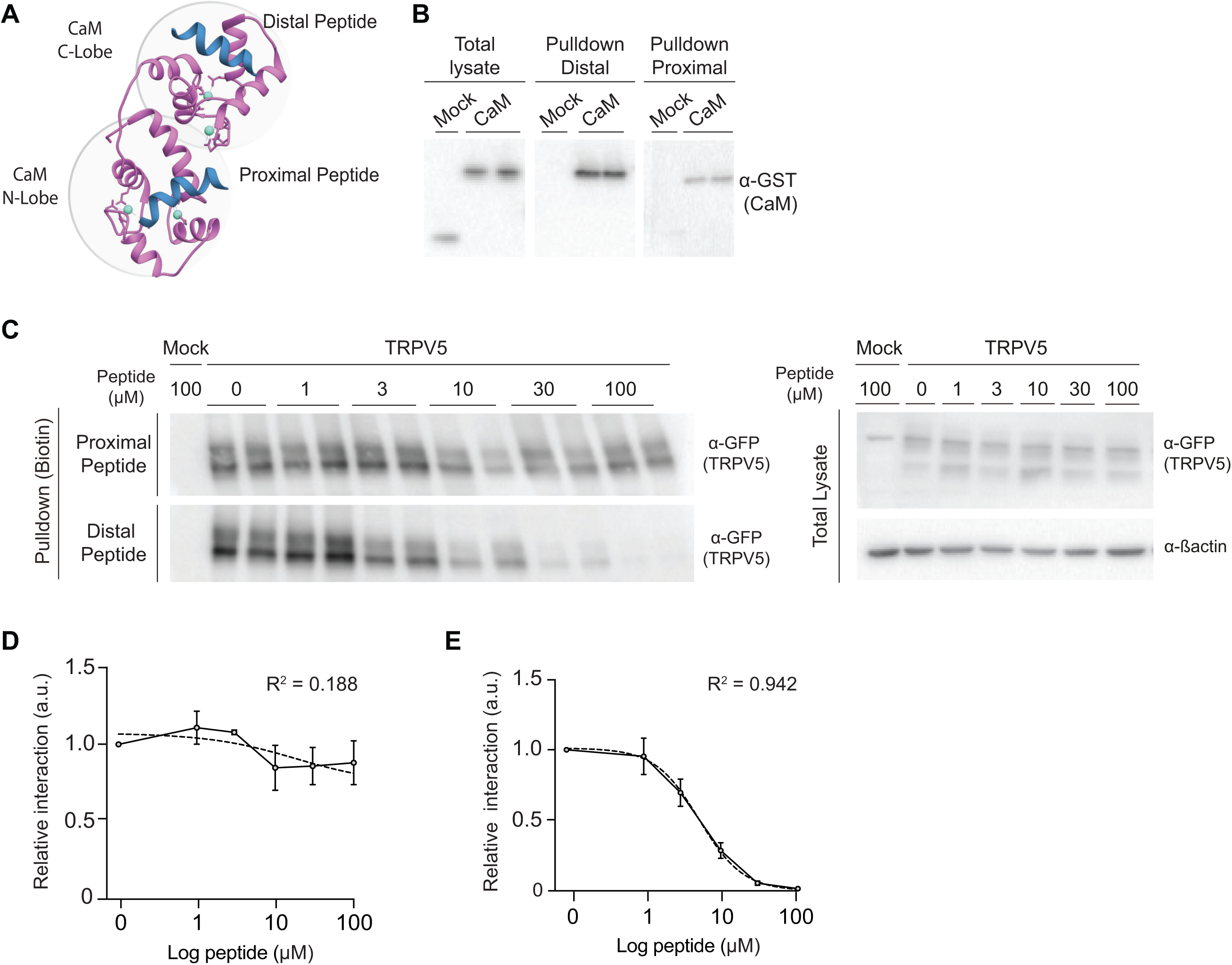
CaM C-lobe as major determinant for binding TRPV5. **A**. Structural overview of the interaction interface of Ca^2+^-bound CaM (pink) with the TRPV5 C-terminal helices (blue). Ca^2+^ ions are depicted as cyan. **B**. Peptide pulldown of mock- or GST-CaM expressing HEK293 cells. Representative immunoblots are shown for total lysate and streptavidin pulldown with either the biotin-linked distal or proximal TRPV5 peptides. **C**. CaM agarose pulldown of HEK293 cells expressing mock eGFP or eGFP-TRPV5, in the presence of increasing amounts (0-100 µM) of either the proximal or distal TRPV5 peptides. Representative immunoblots are shown for eGFP-TRPV5 pulldown and total lysate. **D-E**. Semi-quantification of CaM pulldown experiments expressed as mean ± SEM (n=3). Intensity of TRPV5 pulldown lanes were expressed against the proximal (**D**) or distal (**E**) peptide concentration and the graphs were fitted using a non-linear sigmoidal correlation. R^2^ is indicated in the graph.

Interestingly, single alanine mutations of TRPV5 residues from either the proximal (D90A, F651A) or the distal (W702, W583A) C-terminal interaction interface did not abolish CaM binding as demonstrated by FLIM-FRET analysis (**Figure 2 – figure supplement 2A/B**). CaM binding of TRPV5 D90A and TRPV5 W702A was slightly less compared to TRPV5 wildtype (**Figure 2 – figure supplement 2B**). Given the interaction between CaM1234 and TRPV5 (**Figure 1B**), we explored the potential involvement of these residues in the interaction and found that CaM1234 exhibited the same ability for interacting to all TRPV5 mutants (**Figure 2 – figure supplement 2C**).

### Lobe-dependent interaction in relation to TRPV5 function

To corroborate our interaction studies and evaluate the bilobal role of CaM on TRPV5 activity, we performed Fura-2 Ca^2+^ imaging of HEK293 cells expressing TRPV5 with endogenous CaM, or overexpression with wildtype or mutant CaMs (**Figure 3A**). The TRPV5- expressing cells showed a peak response with plateau after addition of Ca^2+^, which was significantly reduced upon co-expression with wildtype CaM (**Figure 3A/B**). Interestingly, co-expression with CaM12 also resulted in a reduction, suggesting an impaired but remaining ability to inhibit TRPV5. Contrarily, co-expression with CaM1234 or CaM34 yielded an increased Ca^2+^ peak response compared to endogenous CaM (**Figure 3A/B).** This implies that CaM1234 or CaM34 can interfere with endogenous CaM binding via a competitive interaction to TRPV5. As has been suggested by others, it thereby has counteracting effect on endogenous CaM-dependent inhibition (23). This data indicates that the CaM C-lobe Ca^2+^ sensitivity is crucial for TRPV5 inactivation, while the N-lobe does not need to be calcified and may have a support role.

**Figure 3.**
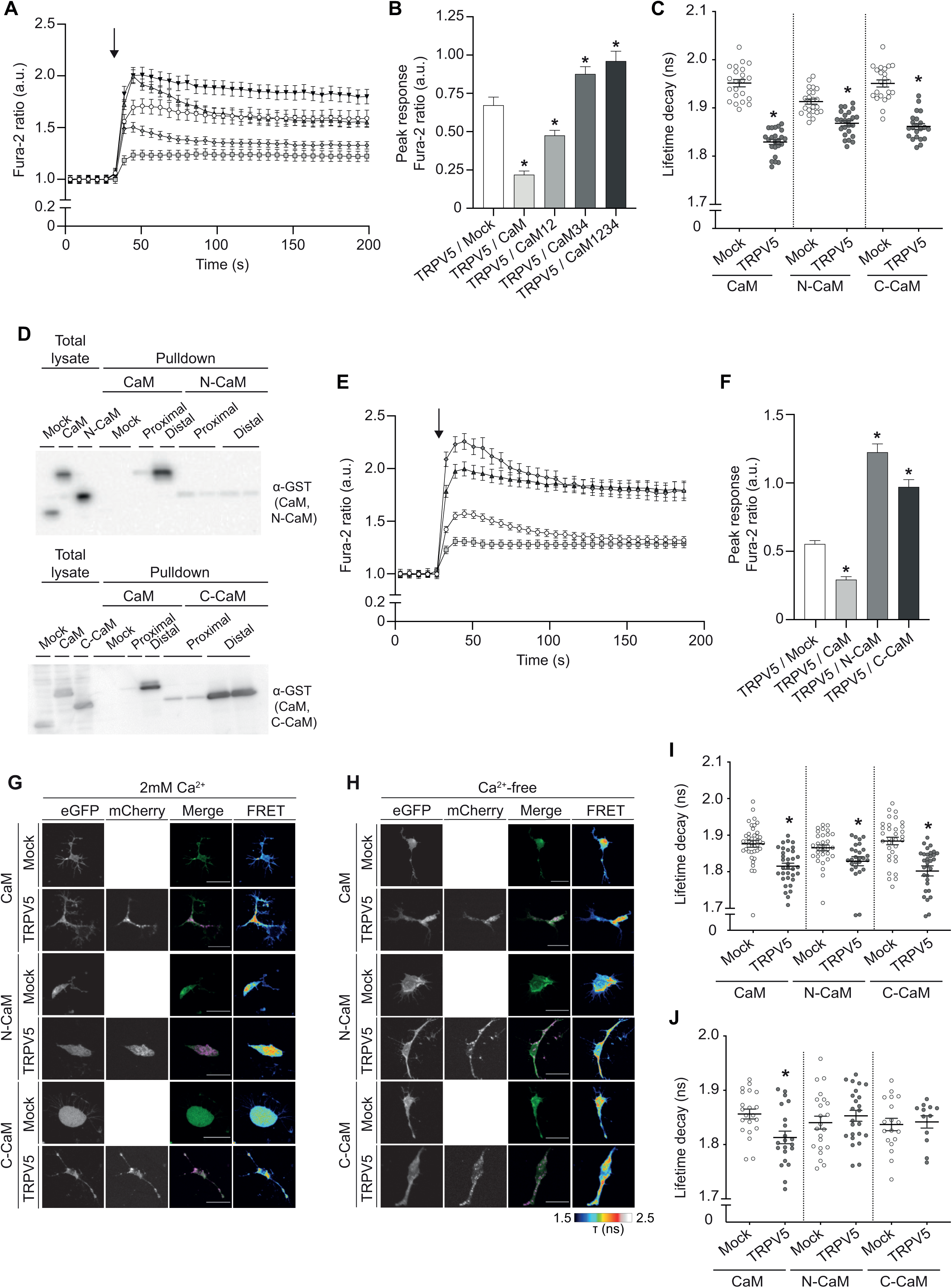
Involvement of the independent CaM lobes in TRPV5 inactivation. **A**. Intracellular Ca^2+^ measurements with Fura2-AM of HEK293 cells expressing mCherry-TRPV5 with either mock or CaM-eGFP wildtype or Ca^2+^-insensitive mutants (CaM12, CaM34, CaM1234). The 340/380nm ratiometric changes are shown as averaged data points ± SEM over time (n=80-160 cells), with addition of Ca^2+^-containing buffer (2 mM Ca^2+^) indicated by the arrow. TRPV5 and mock (open squares), TRPV5 and CaM (light grey circles), TRPV5 and CaM12 (grey upfacing triangles), TRPV5 and CaM34 (dark grey downfacing triangles), TRPV5 and CaM1234 (black diamonds). **B**. Bar graph of the Fura-2 peak response, measured as the ratio of response (mean of four values after Ca^2+^ addition) and baseline (average of five reference values), for the indicated conditions. * indicates p<0.05 compared to TRPV5/mock within each condition (ANOVA). **C**. FLIM-FRET analysis to address the interaction of mCherry-TRPV5 with CaM-eGFP or the separate lobes, CaM N-lobe-eGFP and CaM C-lobe-eGFP, in HEK293 cells. Life time decay of CaM wildtype or N-/C-lobes in the presence of mock or TRPV5, depicted as single measurements of ROIs at cytoplasmic colocalization (n=24-30 cells). * indicates p<0.05 compared to mock within each condition (ANOVA). **D**. Peptide pulldown of HEK293 cells expressing mock, GST-CaM, GST-N-lobe, or GST-C-lobe. Representative immunoblots are shown for total lysate and streptavidin pulldown with either the biotin-linked distal or proximal TRPV5 peptides. **E**. Intracellular Ca^2+^ measurements with Fura2-AM of HEK293 cells expressing mCherry-TRPV5 with either mock, CaM-eGFP wildtype or the separate lobes (CaM N-lobe-eGFP and CaM C-lobe-eGFP). The 340/380nm ratiometric changes are shown as averaged data points ± SEM over time (n=130-200 cells), with addition of Ca^2+^-containing buffer (2 mM Ca^2+^) indicated by the arrow. TRPV5 and mock (open squares), TRPV5 and CaM (light grey circles), TRPV5 and CaM N-lobe (grey diamond), TRPV5 and CaM C-lobe (dark grey upfacing triangles). **F**. Bar graph of the Fura-2 peak response, measured as the ratio of response (mean of four values after Ca^2+^ addition) and baseline (average of five reference values), for the indicated conditions. * indicates p<0.05 compared to TRPV5/mock within each condition (ANOVA). **G-H**. FLIM-FRET analysis of mCherry-TRPV5 with either CaM-eGFP, CaM N-lobe-eGFP or CaM C-lobe-eGFP in PMLs of HEK293 cells washed with Ca^2+^-containing (2 mM) (**G**) or Ca^2+^-free (2 mM EGTA, 2 mM EDTA) buffer (**H**). Representative images are shown for the donors (CaM-eGFP and indicated lobes) and acceptor (mCherry-TRPV5) expression, merged channels (donors: green, acceptor: magenta, co-localization: light pink or white) and FLIM-FRET. Bars represent 10 μm. **I-J**. Life time decay of CaM or N-/C-lobes in the presence of mock or TRPV5, depicted as single measurements of PMLs, for either Ca^2+^-containing (n=25-40 cells) (**I**) or Ca^2+^-free condition (**J**) (n=10-25 cells). ROIs for analysis were drawn throughout the PML and where both proteins colocalized. * indicates p<0.05 compared to mock within each condition (ANOVA). **Figure**

It led us to generate single N- and C-lobes to evaluate whether the lobes can act independently on TRPV5. Following similar steps as abovementioned, we first addressed the interaction capability of both lobes. Either N- or C-lobe of CaM could effectively interact with TRPV5, as shown by the reduction on lifetime decay (**Figure 3C, Figure 3 – figure supplement 3**). The specific interaction of the independent lobes with the TRPV5 helices was studied in a peptide pull-down of HEK293 cells expressing CaM N- or C-lobe (**Figure 3D**). In line with the structural findings of TRPV5-CaM (16, 17), most significant binding was observed between the CaM C-lobe and the distal TRPV5 peptide. Notably, there was also weak but consistent interaction of the N-lobe with both peptides (**Figure 3D**). Next, we addressed the potential independent effect on TRPV5 function. Fura-2 Ca^2+^ imaging demonstrated that co-expression of TRPV5 with either CaM N- or C-lobe increases the intracellular Ca^2+^ peak compared to TRPV5 with endogenous or overexpressed wildtype CaM (**Figure 3E/F**). This indicates that, while both lobes can independently interact with TRPV5, full length CaM is needed for TRPV5 inhibition.

We further explored the interaction of each CaM lobe in Ca^2+^-free conditions using FLIM-FRET analysis of PMLs from cells co-expressing TRPV5 and CaM N- or C-lobe. While CaM and apoCaM interact with TRPV5, the N- and C- lobe interact in Ca^2+^-containing (2 mM), but not in Ca^2+^-free (2 mM EGTA, 2 mM EDTA) conditions (**Figure 3G-J**). Summing up, the C-lobe is the major entity for sensing Ca^2+^ changes and closing TRPV5. The N-lobe (and the linker) act as a support to the structural rearrangement required for the C-lobe to execute its function.

### Stoichiometry of CaM binding to TRPV5

In addition to the dynamics of TRPV5-CaM binding, the stoichiometry of the TRPV5-CaM complex is still under debate. Recent structural work by our group provided evidence for binding of either 1 or 2 CaM molecules (16) to the tetrameric channel, while other groups have described a 1:1 stoichiometry for TRPV5:CaM as well as for TRPV6:CaM (9, 10, 17, 18). To this end, we implemented a method to reveal the *in vivo* stoichiometry of TRPV5-CaM, which is single molecule photobleaching counting (smPB) (24). Following the procedure described in Methods, single molecule complexes at the cell surface were imaged using a TIRF microscope for long acquisition lapses. To optimize the conditions of this technique for our study, we started with HEK293 cells only expressing eGFP-TRPV5 and observed individual molecules at the TIRF plane (**Figure 4A**). Analysis of their traces was performed by filtering them with a tailored Chung-Kennedy Filter Plugin as described previously (25). Some obtained traces matched the expected number of 4 bleaching steps for a tetramer like TRPV5 (**Figure 4B**), while there were also molecules with bleaching steps ranging from 1 to 3. By fitting the observed values to a binomial distribution, we could infer an eGFP folding probability of 72% (**Figure 4C**). Next, HEK293 cells co-expressing mCherry-TRPV5 and eGFP-CaM were subjected to smPB to analyse the stoichiometry of binding. Of note, a slightly different distribution was found for mCherry-TRPV5 compared to eGFP, since the probability of folding was lower (p=0.62, data not shown). In order to assess stoichiometry of binding, low mobile mCherry-TRPV5 spots were chosen that were also positive for eGFP (**Figure 4D**). These were analysed in the presence and absence of extracellular Ca^2+^. The latter condition was taken along to reduce potential changes in intracellular Ca^2+^ levels as a result of Ca^2+^ influx. CaM analysis revealed mainly two type of traces: one or two bleaching steps (**Figure 4E-F**). A highly reduced amount of three steps traces could be found (2% and 5% for presence and absence of Ca^2+^, respectively). Fitting our results to a binomial distribution with eGFP folding probability of 72% showed that the distributions fit with a preferential stoichiometry of 2 CaM molecules per TRPV5 tetramer, either in the presence or absence of extracellular Ca^2+^ (**Figure 4G-H**).

**Figure 4.**
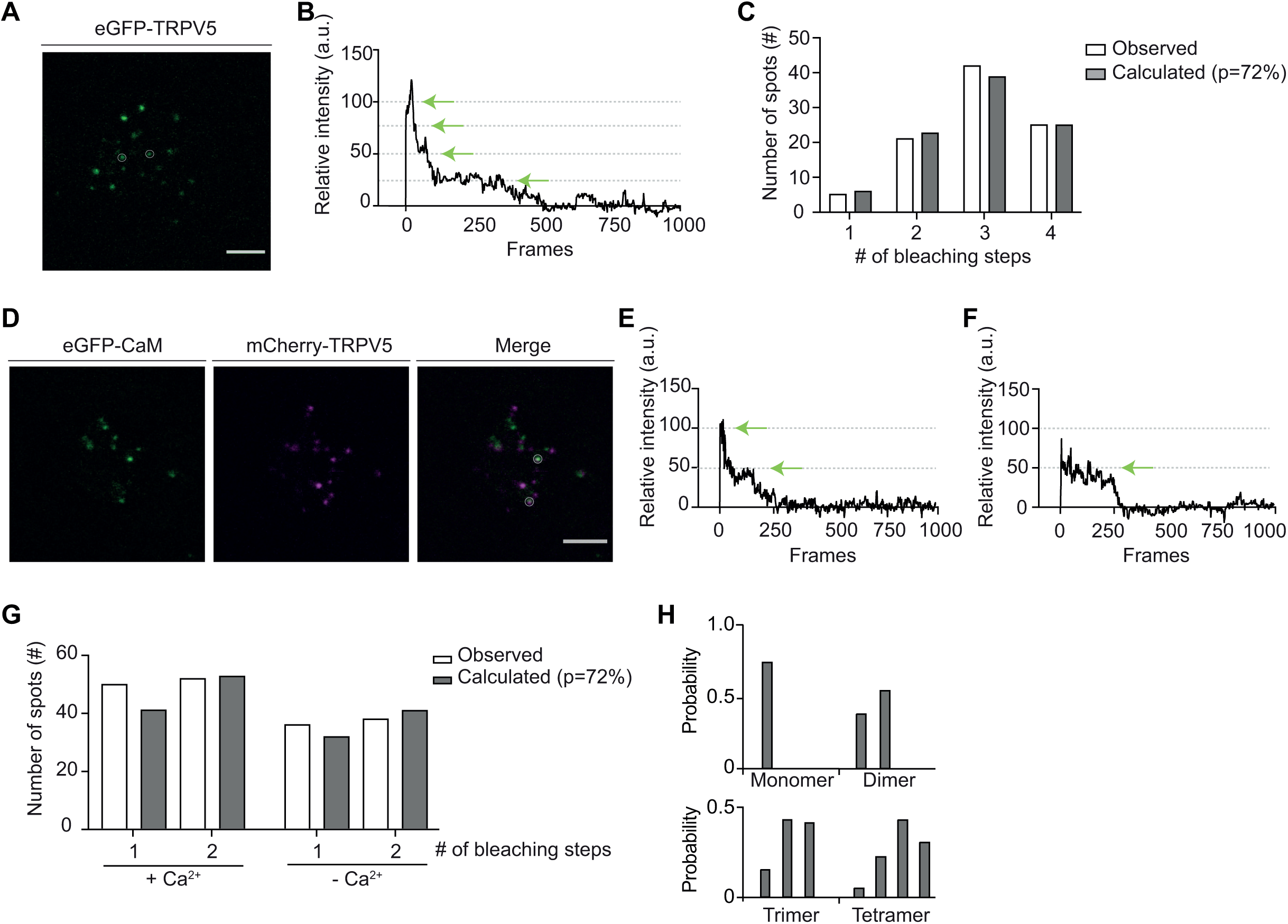
Stoichiometry of CaM binding to TRPV5. **A**. A single frame from a TIRF movie of a HEK293 cell expressing mCherry-TRPV5. White circles indicate immobile spots suitable for counting (scale bar: 5 μm). **B**. Example of four bleaching steps from a spot of mCherry- TRPV5. Dotted lines indicate the fluorescence intensity of single mCherry fluophores. Green arrows point the bleaching steps. **C**. Bar graph with distributions of observed number of bleaching steps (white bars) from eGFP-TRPV5 spots (n=93), and a calculated binomial distribution fit with p (probability of mCherry to be fluorescent) of 72% (grey bars). **D**. Single frames from a TIRF movie of a HEK293 cell expressing mCherry-TRPV5 and CaM-eGFP. White circles indicate immobile spots suitable for counting (scale bar: 5 μm). **E-F**. Examples of bleaching steps from spots of CaM-eGFP, that were positive for mCherry-TRPV5. Dotted lines indicate the fluorescence intensity of single eGFP fluorophores. Green arrows point the bleaching steps. **G**. Bar graph with distributions of observed number of bleaching steps (white bars) from CaM-eGFP, in presence (n=102) and absence (n=74) of Ca^2+^, and a calculated binomial distribution fit with p (probability of eGFP to be fluorescent) of 72% (grey bars). **H**. Theoretical probabilities for monomers, dimers, trimers, and tetramers with p (probability of mEGFP being fluorescent) of 72%.

## Discussion

The present study sheds new light on the dynamic interaction between TRPV5 and CaM and its stoichiometry. It shows a constitutive weak interaction of TRPV5 with apoCaM, and a solid interaction with CaM C-lobe in the presence of 10nM of Ca^2+^, mimicking basal physiological conditions. Importantly, our data provided evidence that a missing CaM N-lobe impacts mildly on the TRPV5 inhibition, and most likely positions the CaM C-lobe. These results establish a new model of CaM-dependent inhibition of TRPV5 in which the C-lobe is responsible for both sensing Ca^2+^ and blocking the channel. Furthermore, we revealed a 1:2 stoichiometry of TRPV5 and CaM in native conditions.

ApoCaM was previously thought of as dormant channel accessory, minimally capable of modulating its targets, but it is now known to trigger a myriad of functionalities across various ion channel families (26). Using various approaches, we now demonstrated for the first-time an interaction between TRPV5 and apoCaM. In contrast to previous studies suggesting that the TRPV5-CaM interaction is only Ca^2+^-dependent (10, 11, 13, 27), our study used the full-length functional proteins in living cells. In this setting, we found that Ca^2+^-dependent inhibition of TRPV5 was counteracted by the presence of the Ca^2+^ - insensitive mutant CaM1234. CaM1234 has often been used to study functional consequences of apoCaM interaction with other channels, with similar dominant-negative effects observed (28-31). It suggests that CaM1234 displaces endogenous CaM from its binding site and thereby prevents channel inhibition. In line with a previous study (32), CaM1234 did not have an effect on the close homologue TRPV6 (data not shown), suggesting that TRPV6 is only regulated by Ca^2+^/CaM. One might speculate that this partly underlies the difference in kinetics of channel inactivation, but further studies need to be conducted to structurally understand this distinction between TRPV5 and TRPV6.

Other examples of pre-association of apoCaM include Ca_v_1.2 and Ca_v_2.1, where it is shown to act as a vigilant sensor regulating channel opening (15). Such regulation is also widely studied in voltage-gated potassium (K_v_) and sodium channels (Na_v_) (33-35). Recent X-Ray crystallography experiments showed the dynamic rearrangement in the Kv7.1, Kv7.4 and Kv7.5 structure of apoCaM and Ca^2+^/CaM (36). Interestingly, TRPV5 is not known to exhibit an IQ calmodulin-binding motif, a feature of many Ca_v_ and Na_v_ channels. Future studies should delineate the exact interaction domain of apoCaM.

An important feature of channel regulation by CaM is the fact that the N- and C-lobes can independently affect channel properties. This is likely due to vastly different affinities (and dissociation kinetics) for Ca^2+^, which in turn can be altered through binding to their targets (26). Once calcified, CaM undergoes a conformational change that likely induces a structural rearrangement within the target proteins to produce a response (26). Generally, the C-lobe presents a higher affinity for Ca^2+^, with the N-lobe requiring higher intracellular Ca^2+^ concentrations to be calcified (37). Free intracellular Ca^2+^ can be variable depending on cellular cell cycle, external insults, amongst other factors. Commonly, mammalian cells are endowed with an average of 100 nM free intracellular Ca^2+^ (38). At this Ca^2+^ concentration, we observed a significant TRPV5-CaM interaction that is likely engaged by a calcified C-lobe, while the N-lobe will remains empty (20). Concomitant with our results, Bokhovchuk and colleagues also described a tight interaction between the CaM C-lobe and the TRPV5 C-terminus at basal Ca^2+^ concentrations (10-100nM) (10). Specifically, we demonstrated that the full-length CaM interaction mainly relies on C-lobe binding to the distal TRPV5 C-terminal helix, as no competition on TRPV5-CaM interaction was observed by the proximal TRPV5 C-terminus peptide. Of note, these distal and proximal peptides are based on helices defined in the cryo-EM TRPV5-CaM structures that were shown to interact with C-lobe and N-lobe, respectively (16, 17). Together, this would be in line with the previously proposed model of CaM-dependent inhibition of TRPV5 (and TRPV6) involving the N-lobe as Ca^2+^ sensor that merely triggers channel closure upon increasing Ca^2+^ concentrations (9, 10, 16, 18, 39). However, our findings suggest that the C-lobe acts as both the sensing and executing unit of CaM. Mutation on the N-lobe (CaM12) did result in CaM-dependent channel inactivation, while an opposite effect was seen for the CaM34 and CaM1234 mutants. The relevance of the C-lobe modulating ion channel activity has been described for other channels, such as of the Kv7 family (Kv7.1, Kv7.4 and Kv7.5) (36).

In addition to these CaM lobe-specific effects, it has been also described for several ion channel families that interaction with both lobes is required to trigger complete Ca^2+^- dependent inhibition (15, 40). This is likely due to a structural rearrangement of the full complex. Indeed, our experiments with single CaM lobes revealed that, while the calcified C- lobe can interact with the channel, it requires an additional part of the CaM structure to fully trigger full Ca^2+^-dependent inactivation of TRPV5. Therefore, we propose a calmodulation model in which TRPV5 and apoCaM are pre-associated (**Figure 5, left panel**) to induce fast, yet non-persistent, closure of the lower channel gate upon calcification of the C-lobe of CaM at locally increased intracellular Ca^2+^ concentrations (**Figure 5, central panel**). Upon a further raise in the intracellular Ca^2+^ concentration, the N-lobe will become calcified and acts as stabiliser of the closed conformation (**Figure 5, right panel**). Upon decrease of local Ca^2+^ (i.e. as result of Ca^2+^ shuttling proteins such as calbindin-D_28K_), the complex rearranges and the CaM C-lobe will open the bottom gate.

**Figure 5.**
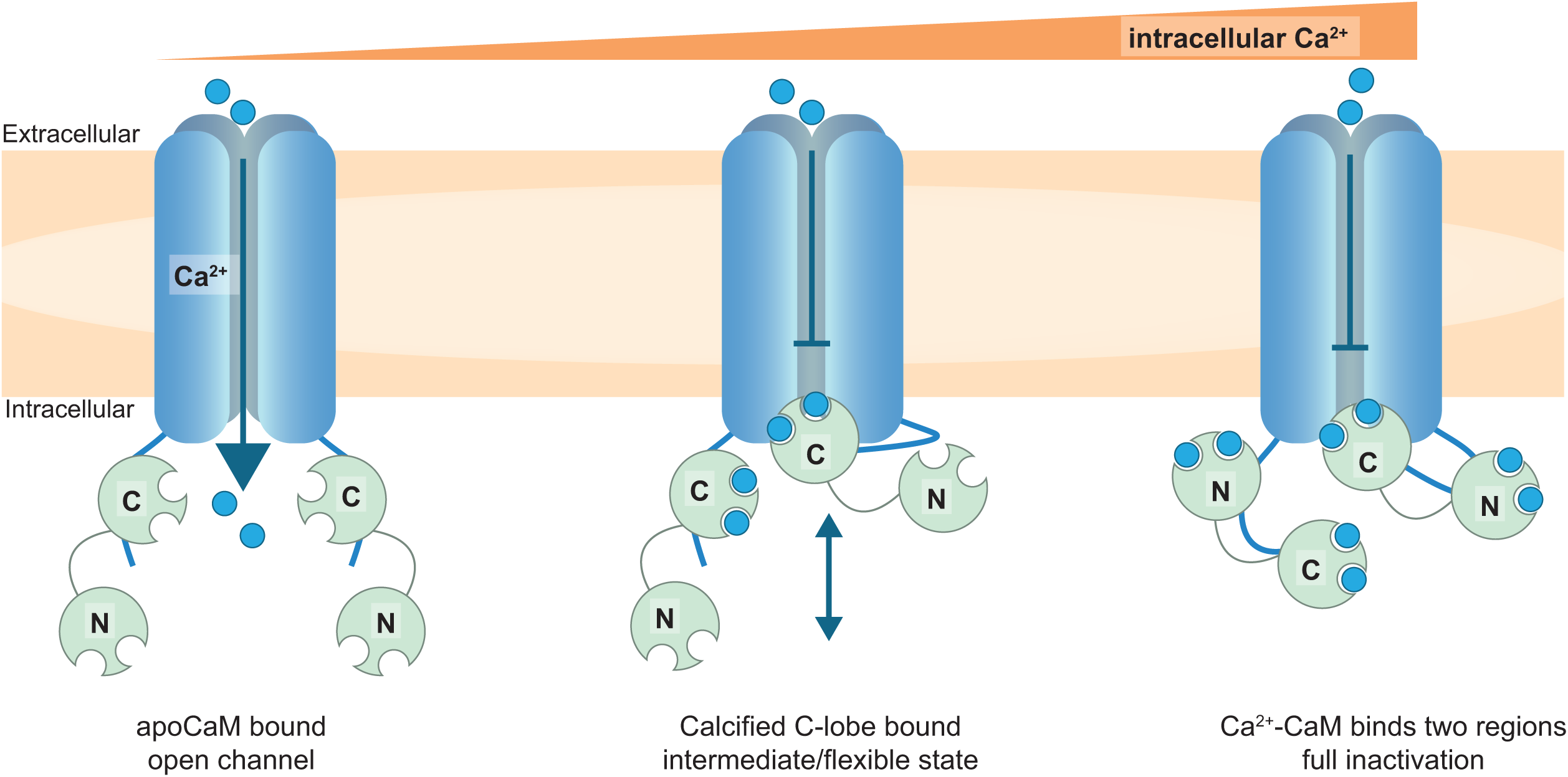
Schematic model of TRPV5 calmodulation. ApoCaM pre-associates with TRPV5, but it is not blocking ion conduction of the channel (left panel). At low basal Ca^2+^ concentrations, the C-lobe can get calcified and obstruct the bottom pore of TRPV5 (central panel). Upon increased Ca^2+^ concentrations, the N-lobe will interact with Ca^2+^ and further enhance the inactivated complex (right panel). There are 2 CaM molecules that can interact with a TRPV5 channel at any given time, while there is only 1 CaM C-lobe effectively blocking the channel pore.

Next to the lobe-specific effects, this study sheds new light on the stoichiometry of binding. We implemented the smPB technique to decipher the amount of CaM molecules per TRPV5 tetramer (41), and identified a preferable 1:2 TRPV5-CaM composition. So far, there has been debate about the stoichiometry. Initial NMR studies with peptides of TRPV5 C-terminus suggested a 1:2 stoichiometry for CaM:TRPV5^696-729^ (12). Later studies of the same group, using longer TRPV5 C-terminus peptides, confirmed these results (10). Yet, recent cryo-EM complex structures of TRPV5 with CaM suggest on one hand a 1:1 stoichiometry of 1 tetrameric TRPV5 channel binding to 1 CaM molecule (17), and on the other hand provide evidence for a variable stoichiometry of either 1:1 or 1:2 (2 CaM molecules per TRPV5 tetramer) (16). Through the smPB method, we were the first to reveal in intact cells that TRPV5 preferably binds 2 CaM molecules.

Since some ion channels undergo changes in stoichiometry depending on the Ca^2+^ concentration, experiments were performed in the presence and absence of Ca^2+^ (42). For example, the stoichiometry of L-type voltage-gated Ca^2+^ channels was found to be 1:1 in the absence of intracellular Ca^2+^, but increased to two CaM peptides at higher Ca^2+^ concentrations (43). We observed the same stoichiometry in the presence and absence of extracellular Ca^2+^, suggesting that basal interaction with CaM is not dependent on Ca^2+^ influx. Analyzing our cryo-EM TRPV5-CaM complex structure (16), it is clear that only 1 CaM molecule can occupy the lower cavity of the tetrameric channel pore. Therefore, we would speculate that the CaM-mediated TRPV5 inactivation mimics the ball-and-chain mechanism observed between Kv channels and Kvβ subunits (44).

In conclusion, our study demonstrates that the CaM-mediated TRPV5 channel inactivation is a dynamic process involving 2 CaM molecules in close proximity to the channel pore that exhibit lobe-specific regulation. The results established a weak, but persistent, interaction between TRPV5 and apoCaM that may assist fast channel inhibition through calcification of the CaM C-lobe in close vicinity of the channel. Hereby, the N-lobe serves primarily as stabiliser of the inactivated TRPV5-CaM complex.

## Materials and Methods

### Molecular cloning

Restriction enzyme digestions, DNA ligations and other recombinant DNA procedures were performed using standard protocols. Rabbit TRPV5 was cloned into a pcDNA5 vector containing a N-terminal GFP (green fluorescent protein) tag for biochemical assays, and a pmCherry vector containing a N-terminal mCherry tag for microscopy experiments. Rat CaM and truncated forms of the protein, N-lobe (aa1–80) and C-lobe (aa81–149), were cloned into a peGFP vector containing a N-terminal eGFP tag for microscopy, and a pEBG vector containing a N-terminal GST tag for biochemical studies. Mutagenesis was performed to generate the indicated CaM mutants, using the QuikChange site-directed mutagenesis method (Stratagene, San Diego, USA) following manufacturer’s protocol. The (partial) Ca^2+^- insensitive mutants CaM12, CaM34, and CaM1234 were based on mutations (D → A) in the EF-hand structures of the N- and C-lobes, known to affect Ca^2+^-binding (45, 46). All DNA constructs were verified by DNA sequencing. DNA for mammalian cell transfection was amplified in *E. coli* TOP10f strain and plasmid preparation was done using the Macherey-Nagel^TM^ Nucleobond^TM^ Xtra Midi kit according to manufacturers’ protocol.

### Buffers

Lysis buffer: 50 mM Tris-HCl (pH 7.5), 150 mM NaCl, 2 mM EGTA, 1% (v/v) Triton X-100, 1 mM sodium orthovanadate, 10 mM sodium-glycerophosphate, 50 mM sodium fluoride, 10 mM sodium pyrophosphate, 0.27 M sucrose, and the freshly added protease inhibitors pepstatin A (1 μg/ml), PMSF (1 mM), leupeptin (5 μg/ml), and aprotinin (1 μg/ml). TBS-Tween (TBS-T): Tris-HCL (200 mM, pH 7.5), 150 mM NaCl, and 0.2% (v/v) Tween-20. Leammli sample buffer (5x): 10% (w/v) SDS, 25% (v/v) β-mercapto-ethanol, 50% (v/v) glycerol, 0.3 M Tris-HCl (pH 6.8), 0.05% (v/v) bromophenol blue. Fura-2 wash buffer: 132.0 mM NaCl, 4.2 mM KCl, 5.5 mM D-glucose, 10 mm HEPES/Tris, pH 7.4. Fura-2 Ca^2+^ buffer: 132.0 mM NaCl, 4.2 mM KCl, 1.4 mM CaCl_2_, 1.0 mM MgCl_2_, 5.5 mM D-glucose, 10 mm HEPES/Tris, pH 7.4. Fura-2 Ethylenediaminetetraacetic acid (EDTA) buffer: 132.0 mM NaCl, 4.2 mM KCl, 2 mM EDTA, 5.5 mM D-glucose, 10 mm HEPES/Tris, pH 7.4. Plasma membrane lawn preparation: 70 mM KCl and 30 mM HEPES pH 7.5, adjusted with KOH (KH buffer).

### Cell culture

HEK293 (human embryonic kidney 293) cells were grown in Dulbeccos Modified Eagles Medium (DMEM, Lonza, Basel, Switzerland) supplemented with 10% (v/v) fetal bovine serum (BioWest, Nuaillé, France), 2 mM L-glutamine, and 10 μl/ml non-essential amino acids at 37°C in a humidity-controlled incubator with 5% (v/v) CO_2_. For transient transfection, cells were transfected 6-8 hours after seeding with the respective DNA construct using polyethyleneimine (PEI, Brunschwig Chemie, Basel, Switzerland) with a DNA:PEI ratio of 1:6. The cells were cultured for 16-36 additional hours prior to the experiments.

### Antibodies

GFP antibody (1:5,000; G1544), GST antibody (1:5,000; G7781), and beta-actin (1:10,000; A5441) were purchased from Sigma Aldrich (Sant Luis, USA). Secondary antibodies coupled to horseradish peroxidase used for immunoblotting were obtained from Sigma (A4914 Goat anti-Rabbit IgG).

### Plasma membrane lawns preparation

Plasma membrane lawns (PML) are cell membrane sheets obtained by unroofing cells with an osmotic shock (47, 48). In short, HEK293 cells were seeded on 18 mm diameter round coverslips coated with Poly-L-Lysine (PLL) and transfected with the respective plasmids. DNA amounts of 650 ng and 1350 ng were used for transfection of donor and acceptor DNA plasmids respectively. After 24 hours of transfection, cells were washed once with ice-cold PBS. Membrane labelling was performed by incubating cells 10 minutes with Wheat Germ Agglutinin Alexa Fluor^TM^ 680 Conjugate (WGA-AF680, Thermo Fisher Scientific, Waltham, USA) diluted 1:100 in DMEM:30 mM HEPES. Next, two washes of 5 minutes with ice-cold PBS were performed to remove unbound WGA-AF680. The osmotic shock was done by incubating cells for 5 minutes with ice-cold KH buffer (diluted 3 times), followed by a gentle wash with non-diluted KH buffer. In order to control the Ca^2+^ content, buffer modifications were incorporated. In Ca^2+^ experimental conditions, KH buffer was supplemented with 2 mM CaCl_2_, while 2 mM EGTA and 2 mM EDTA were added for the Ca^2+^-free conditions. After two washes of 5 minutes with the respective ice-cold KH buffer, only unroofed cells remain attached. Their membranes were fixed with fresh 4% (v/v) paraformaldehyde for 10 minutes at room temperature and mounted in Fluoromont-G^TM^ mounting media (Thermo Fisher Scientific, Waltham, USA).

### Co-immunoprecipitation assay

HEK293 cells expressing GFP-TRPV5 and GST-tagged CaM wildtype or mutants were lysed (lysis buffer containing 5 mM CaCl_2_) 36 hours after transfection. Lysates were cleared by centrifugation at 4°C for 15 minutes at 16,000 g and protein concentration was measured by the Bradford method (Bio-Rad, Hercules, USA). Next, 1 mg lysate was incubated with glutathione agarose resin (GE Healthcare, Chicago, USA) for 2 hours at 4**°**C under gentle rotation. Following three washes with lysis buffer (5 mM CaCl_2_), proteins were eluted in 30 μl of 2X Laemmli sample buffer.

### Peptide pull down assays

The following peptides were purchased from EMC microcollections GmbH: N’-Biotin-C6-QSSNSHRGWEILRRNTLGHL-C’ (TRPV5 distal helix), N’-Biotin-C6-QSSNSHRGAEILRRNTLGHL (W702A), and Biotin-C6-ENHHDQNPLRVLRYVEAFKCSDKEDGQ-C’ (TRPV5 proximal helix). For a peptide pull down, HEK293 cells expressing GST-tagged CaM or CaM lobes were lysed 36 hours after transfection. Lysis buffer contained 5 mM CaCl_2_. 1 mg of cleared lysate was incubated with 3 μg of the respective peptides for 10 minutes at 4°C, followed by 5 minutes incubation with 20 μl streptavidin agarose resin (Thermo Fisher Scientific, Waltham, USA) at 4°C under gentle rotation. The samples were washed three times with lysis buffer (5 mM CaCl_2_) and proteins were eluted in 30 μl of 2X Laemmli sample buffer.

### CaM binding assay

HEK293 cells expressing GFP-TRPV5 were lysed 36 hours after transfection, in buffers containing different Ca^2+^ concentrations as indicated. These were prepared by adding CaCl_2_ to the lysis buffer (containing 2 mM EGTA) in order to obtain the indicated free Ca^2+^ concentrations, as calculated using MaxChelator (https://somapp.ucdmc.ucdavis.edu/pharmacology/bers/maxchelator/CaEGTA-TS.htm). Next, 1 mg of cleared lysate was incubated with CaM agarose resin (Sigma Aldrich, San Luis, USA) for 2 hours at 4**°**C under gentle rotation. In case of peptide competition, 0-100 µM peptide was added during this period. The immunoprecipitates were subsequently washed three times with lysis buffer containing the indicated CaCl_2_ concentration, and proteins were eluted in 30 μl of 2X Laemmli sample buffer.

### Immunoblotting

All samples (immunoprecipitates and total cell lysates) were subjected to 8-12% (w/v) SDS-PAGE and transferred to polyvinylidene fluoride (PDVF) membranes. These membranes were blocked for 30 minutes with 5% (w/v) non-fat dry milk (NFDM; in TBS-T) and immunoblotted overnight at 4°C using indicated primary antibodies. Next, the blots were washed with TBS-T, incubated with secondary peroxidase-labelled secondary antibodies (in 5% (w/v) NFDM/TBS-T) for 1 hour at room temperature. Following repeated washes, protein expression was visualized with chemiluminescence SuperSignal West reagent (Thermo Fisher Scientific, Waltham, USA) using the Bio-Rad ChemiDoc XRS imaging system.

#### FLIM-FRET (Förster Resonance Energy Transfer with Fluorescence Lifetime Imaging Microscopy)

HEK293 cells were seeded on PLL coated coverslips (18 mm diameter) and transfected with indicated donor (450 ng, eGFP) and acceptor (950 ng, mCherry) plasmids, and/or corresponding mock DNA for 24 hours. For PML, check *Plasma membrane lawns preparation section*.

Time-domain FLIM was performed with a 63X/1.2 numerical aperture (NA) Plan Apochromat water objective lens on a Leica SP8 SMD laser scanning confocal system. Samples were excited at 488 nm, 10% laser power, by a high energy pulsed IR-fiber white laser with pulses of 200 ps at 80 MHz. Fluorescence photons were detected with hybrid detectors (495 nm – 545 nm detection range) in photon counting mode using a TCSPC approach operated by the FALCON module (Leica Microsystems, Manheim, Germany) integrated within the Leica SP8 SMD system. Cells with at least 100-1,000 photons per pixel were acquired and incorporated to the analysis. FLIM module on LASX was utilized to acquire the fluorescence decay of Regions of Interest (ROI) drawn on extranuclear donor-acceptor colocalization areas of the cell or PMLs (FALCON Leica software, Wetzlar, Germany). The *n-Exponential Reconvolution* fitting model was implemented for mono-exponential decay fitting, which also deconvoluted the instrument response function (IRF).

### Single Molecule Photobleaching (smPB)

HEK293 cells were seeded on 12-well dishes and transfected with low amounts of the DNA plasmids of interest for 16 hours (100 ng TRPV5, 150 ng of CaM). Parallelly, bottom glass Willco-dishes® (WillCo Wells B.V, Amsterdam, the Netherlands) were incubated with Concanavalin A (2mg/ml) diluted in PBS for 30 minutes at 37°C and washed O/N with PBS at the same temperate. Next, cells were resuspended with Trypsin (Lonza, Basel, Switzerland) and reseeded with fresh DMEM media (check *Cell culture* section) on the Willco-dishes previously prepared. Imaging was performed from 30 minutes to 5 hours post-reseeding.

Total internal reflection microscopy (TIRFM) was used for illuminating complexes located at the cell surface. To this end, we used a 150X/1.45 plan apochromat oil objective on an Olympus IX-71 widefield fluorescence microscope equipped with a TIRF system (Olympus, Shinjuku, Japan) and an EM-CCD camera (Hamamatsu ImagEM, Hamamatsu Photonics, Hamamatsu, Japan). eGFP and mCherry were excited with a 488 nm Argon laser and a 594 nm DPSS laser. Movies of 2,000 frames of a 54 x 54 um^2^ area were acquired at 30 frames per second.

First, fluorescence from mCherry was recorded under illumination with 593 nm light. After most of mCherry fluorescent spots were bleached out, 593 nm light was switched off and subsequently 488 nm light turned on, and mEGFP fluorescence was recorded. Spots that stayed confined throughout the whole movie were selected for counting.

Movies were opened in Fiji (49) and confined spots were selected. To filter out the noisy traces obtained, we developed a Fiji plug-in based on the NoRSE algorithm (50), which adapted and optimised the Chung-Kennedy filter (51). Previous studies followed a similar procedure to clearly reveal the bleaching steps (25, 52, 53). ROIs were drawn to include the confined spots no bigger than 10 pixels^2^. Traces of confined spots and background areas were filtered with a sharpness factor (p) of 40, a change detection range (N) of 20 and 4 sample windows with sizes (M) of 4, 8, 16 and 32. Background traces were mathematically subtracted from spot traces and intensity decays were manually counted.

### Intracellular Ca^2+^ measurements using Fura-2-AM

HEK293 cells expressing mCherry-TRPV5 and eGFP-CaM wildtype or mutants were seeded in fibronectin-coated Press-to-Seal silicone isolator wells (Molecular Probes; diameter of 2.5 mm) on Superfrost Plus Microscope Slides (Thermo Fisher Scientific, Waltham, USA). After 2-4 hours, cells were loaded with 3 μM Fura-2-AM and 0.01% (v/v) Pluronic F-129 (both from Molecular Probes) in DMEM medium at 37 °C for 20 minutes. After loading, the cells were washed twice with Fura-2 wash buffer and allowed to equilibrate for another 10 minutes in Fura-2 EDTA buffer. Next, the slide was placed on an inverted microscope (Axiovert 200M, Carl Zeiss, Jena, Germany) at 20X magnification, and intracellular Ca^2+^ levels were monitored after fluorescence excitation at 340 and 380 nm using a monochromator (Polychrome IV, TILL Photonics, Gräfelfing, Germany). Fura-2 buffer containing CaCl_2_ was added after reaching a steady basal state. Fluorescence emission light was directed by a 415DCLP dichroic mirror (Omega Optical, Inc., Brattleboro, VT) through a 510WB40 emission filter (Omega Optical Inc. Brattelboro, USA) onto a CoolSNAP HQ monochrome CCD-camera (Roper Scientific, Vianen, the Netherlands). The integration time of the CCD-camera was set at 200 milliseconds with a sampling interval of 3 seconds. All hardware was controlled with Metafluor (version 6.0) software (Universal Imaging Corp., Downingtown, PA). For each wavelength, the mean fluorescence intensity was monitored in an intracellular region and, for purpose of background correction, an extracellular region of identical size, both set prior to the start of the experiment. After background correction, the fluorescence emission ratio of 340 and 380 nm excitation was calculated to determine changes in intracellular Ca^2+^ concentration. The peak response is calculated as the difference in ratio upon Fura-2 Ca^2+^ buffer addition versus basal levels in Fura-2 EDTA buffer (t_Ca_ - t_EDTA_). All measurements were performed at room temperature.

### Statistical analysis

The immunoblot data were analyzed by comparing integrated optical densities of bands using Fiji (49). The semi-quantification is shown as mean ± SEM and plotted against the log free Ca^2+^ concentration (Figure 1E) or log peptide concentration (Figure 2). FLIM-FRET data is shown as individual data points with a mean ± SEM indicated in each graph, with n as the number of cells. The Fura-2-AM measurements are depicted as averaged 340/380 ratio ± SEM over time, with n as the number of cells, and a bar graph showing the peak response (t_Ca_ - t_EDTA_) as mean ± SEM. For all data, p < 0.05 was considered statistically significant using a one-way ANOVA with a Dunnett’s multiple comparison posthoc test. Data representation and analysis was done using Fiji software (49) and GraphPad Prism 8.0 (GraphPad Software Inc., San Diego, USA). For the Fura-2-AM data, Graphpad was used to test for normality of distribution and identify outliers. In case of non-normal distribution, a Kruskal-Wallis test with Dunn’s multiple comparisons test was performed.

## Acknowledgements

We thank Mark van Goor for helpful suggestions and discussions. SRR was supported by *Alfonso Martín Escudero* Grant for Postdoctoral studies abroad. This study was financially supported by the Netherlands Organization for Scientific Research (VICI 016.130.668) and EU Horizon 2020 Marie Skłodowska-Curie Actions (748058).

## Competing interests

The authors declare no conflict of interest

## Author contributions

SRR and JvW conceived and designed the experiments. SRR and NT performed the experiments. MvE developed the Fiji Plugin for the smPB. SRR, NT and JvW analysed the data. SRR and JvW wrote the manuscript, with approval of all authors. JF, JGH and JvW supervised the work.

## Figure legends

**Figure 1 – figure supplement 1.**
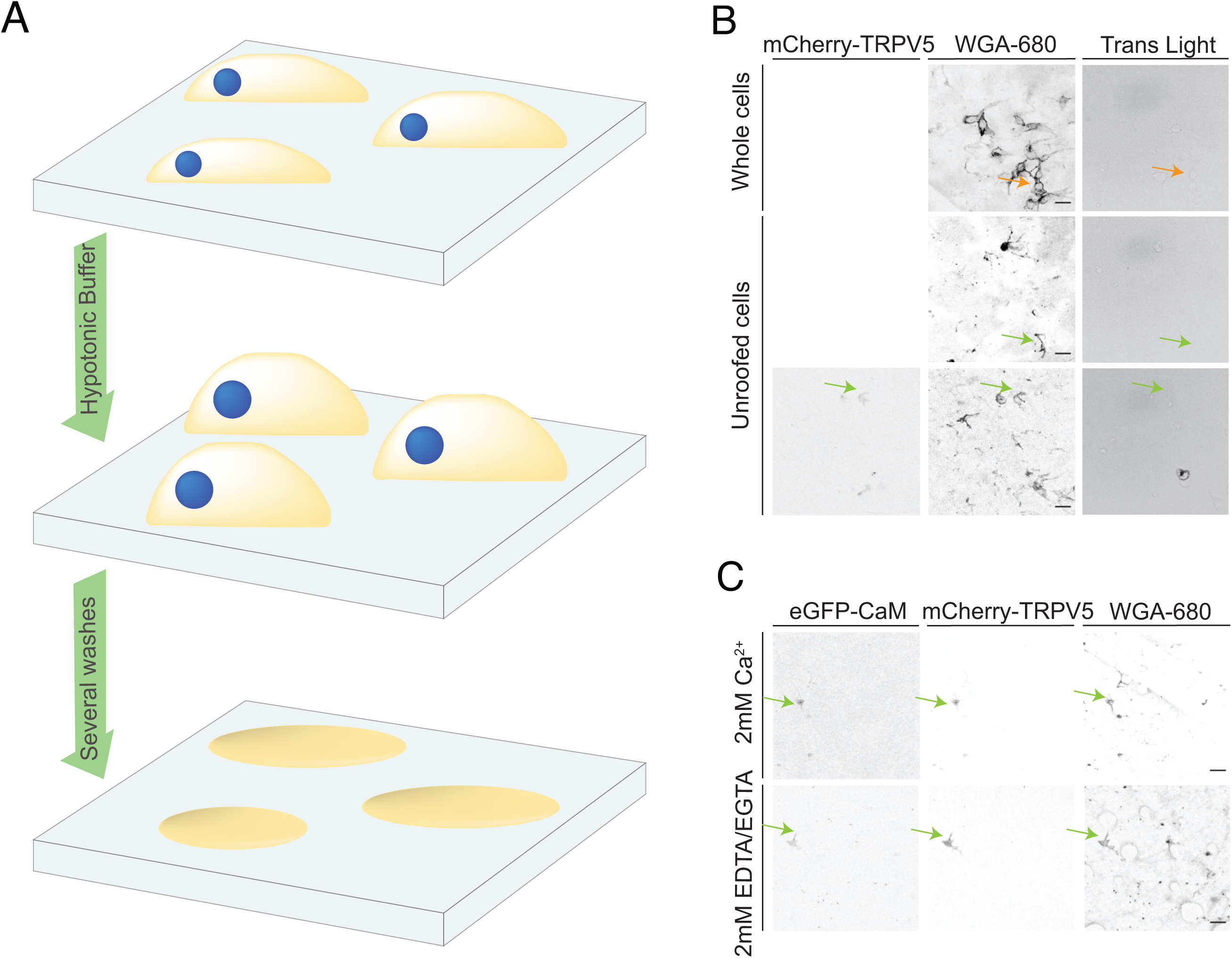
Preparation of plasma membrane lawns (PMLs). **A**. Schematic overview of the protocol setup. HEK293 cells were washed with hypotonic buffer for cell swelling, followed by sequential washes with the Ca^2+^-containing or Ca^2+^-free buffer that lead unroofing the cells leaving the membranes attached to the glass coverslip. **B**. Representative images are shown for fluorescent imaging of mCherry-TRPV5 and the WGA- 680 membrane marker as well as of transmitted light in the whole cells and unroofed cells. Bars represent 20 μm. **C**. Representative images of eGFP-CaM, mCherry-TRPV5 and the WGA-680 expressed or labelled on unroofed cells. Bars represent 20 μm. Orange arrows point to whole cells, green arrows to PMLs.

**Figure 1 – Source data 1. Original immunoblot density calculations for figure 1E**

**Figure 2 – figure supplement 1.**
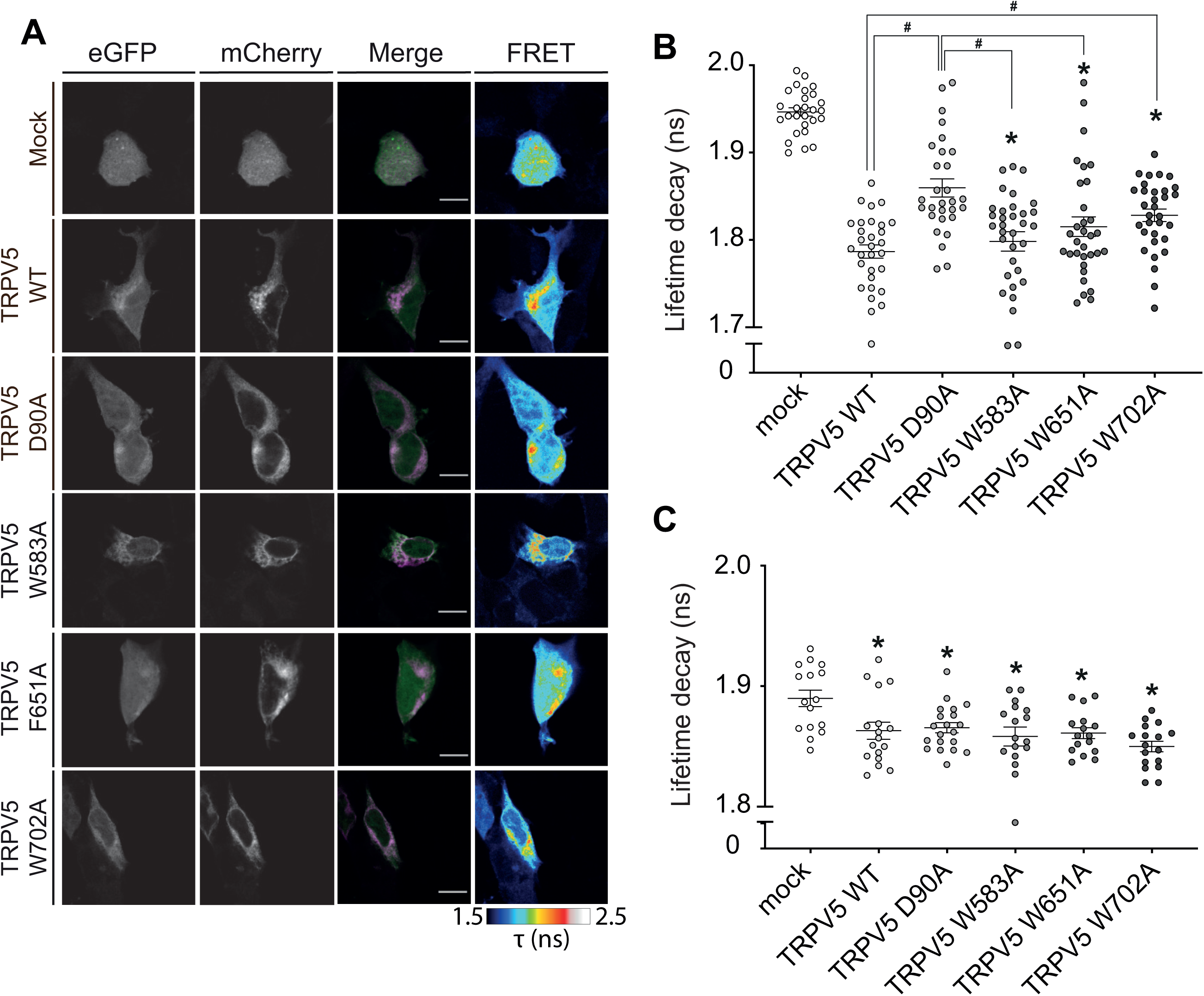
Interaction interface of TRPV5-CaM. Interaction of mCherry-TRPV5 wildtype and indicated mutants with CaM-eGFP in HEK293 cells was addressed by FLIM-FRET. **A.** Representative images are shown for the donor (CaM-eGFP) and acceptors (mCherry-TRPV5 WT and mutants) expression, merged channels (donors: green, acceptor: magenta, co-localization: light pink or white) and FLIM-FRET image. Scale bars represent 10μm. **B**. Life time decay of CaM in the presence of mock, TRPV5 wildtype or indicated mutants, depicted as single measurements of ROIs at cytoplasmic colocalization (n=25-30 cells). **C**. Life time decay of CaM1234 in the presence of mock, TRPV5 wildtype or indicated mutants, depicted as single measurements of ROIs at cytoplasmic colocalization (n=15-20 cells). * indicates p<0.05 compared to mock (ANOVA), and # indicates p<0.05 compared to the TRPV5 wildtype condition (ANOVA).

**Figure 2 – Source data 1. Original immunoblot density calculations for figure 2D**.

**Figure 2 – Source data 2. Original immunoblot density calculations for figure 2E.**

**3 – figure supplement 1.**
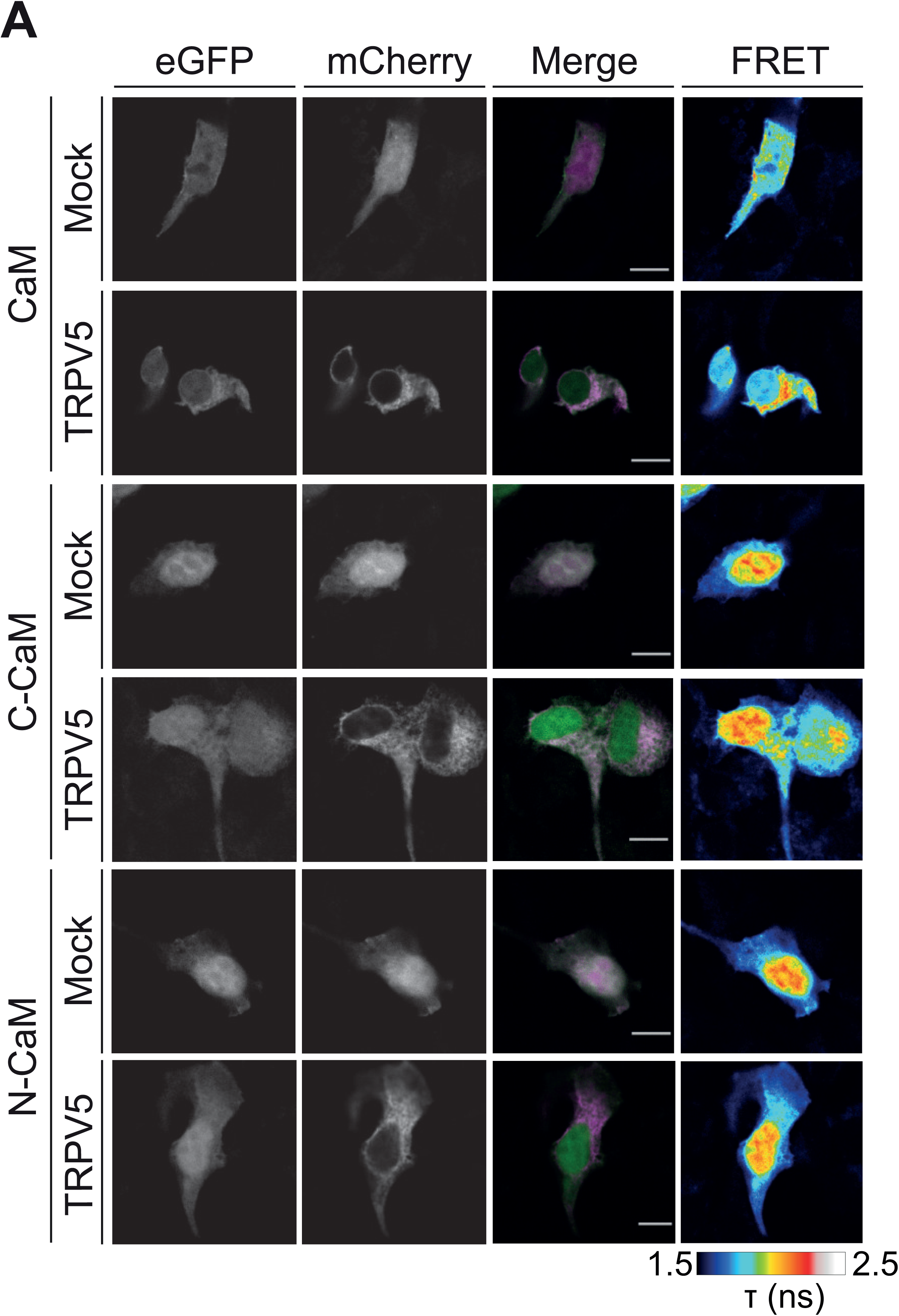
N- and C-Lobe interaction with TRPV5. Interaction of CaM WT, N-Lobe and C-Lobe with mCherry-TRPV5 in HEK293 cells was addressed by FLIM-FRET. **A.** Representative images are shown for donors (CaM, N-CaM and C-CaM-eGFP) and acceptors (mock or mCherry-TRPV5) expression, merged channels (donors: green, acceptor: magenta, co-localization: light pink or white) and FLIM-FRET image. Scale bars represent 10 μm.

**Figure 3 – Source data 1. Original peak response values for figure 3B**.

**Figure 3 – Source data 2. Original peak response values for figure 3F**.

**Figure 4-source data 1. Original values on observed and calculated number of spots for figure 4C**.

**Figure 4-source data 2. Original values on observed and calculated number of spots for figure 4G**.

**Figure 4-source data 3. Original values on probabilities for figure 4H**.

